# Seasonal plasticity in anti-predatory strategies: matching of color and color preference for effective crypsis

**DOI:** 10.1101/253906

**Authors:** Erik van Bergen, Patrícia Beldade

**Affiliations:** Instituto Gulbenkian de Ciência, Oeiras, Portugal; Research Centre for Ecological Change, Faculty of Biological and Environmental Sciences, University of Helsinki, Helsinki, Finland; UMR5174 - CNRS, Evolution et Diversité Biologique, Université Paul Sabatier, Toulouse, France

**Keywords:** animal coloration, background matching, seasonal environments, phenotypic plasticity

## Abstract

Effective anti-predatory strategies typically require matching appearance and behavior in prey, and there are many compelling examples of behavioral repertoires that enhance the effectiveness of morphological defenses. When protective adult morphology is induced by developmental environmental conditions predictive of future predation risk, adult behavior should be adjusted accordingly to maximize predator avoidance. While behavior is typically strongly affected by the adult environment, developmental plasticity in adult behavior — mediated by the same pre-adult environmental cues that affect morphology — could ensure an effective match between anti-predatory morphology and behavior. The coordination of environmentally-induced responses may be especially important in populations exposed to predictable environmental fluctuations (e.g. seasonality). Here, we studied early and late life environmental effects on a suite of traits expected to work together for effective crypsis. We focused on wing color and background color preference in *Bicyclus anynana*, a model of developmental plasticity that relies on crypsis as a seasonal strategy for predator avoidance. Using a full-factorial design, we disentangled effects of developmental and adult ambient temperature on both appearance and behavior. We showed that developmental conditions affect both adult color and color preference, with temperatures that simulate natural dry season conditions leading to browner butterflies with a perching preference for brown backgrounds. This effect was stronger in females, especially when butterflies were tested at lower ambient temperatures. In contrast to the expectation that motionlessness enhances crypsis, we found no support for our hypothesis that the browner dry-season butterflies would be less active. We argue that the integration of developmental plasticity for morphological and behavioral traits might improve the effectiveness of seasonal anti-predatory strategies.

IMPACT SUMMARY

To avoid predation, prey rely on strategies that typically include a variety of morphological and behavioral characteristics working together to deceive or scare predators. While some protective traits are a constitutive property of the prey species (e.g. hedgehog spines), others are produced only when the risk of encountering predators is high. For example, *Daphnia* crustaceans develop protective ‘helmets’ and spines when exposed to cues that signal predator presence. When anti-predator morphologies are environmentally-induced, or plastic, the responses of associated behavioral traits should also be environmentally-dependent to ensure that animals exhibit behavioral repertoires that match their appearance.

We used the tropical butterfly *Bicyclus anynana*, the squinting bush brown, to study these coordinated responses. In its natural habitat, with alternating dry and wet seasons, this thermally plastic butterfly alternates between seasonal forms with distinct wing color patterns related to distinct strategies to avoid predation. Dry season individuals, which develop under cooler temperatures, have small pattern elements on their wings and are cryptic against the brown background of dry foliage. In contrast, wet season individuals have large ornamental pattern elements that deflect attacks from predators towards the wing margin and away from their vulnerable bodies.

We tested whether the cooler temperatures of the dry season also influenced other traits that can presumably improve the effectiveness of camouflage. In particular, we found that both the ornamental colors (appearance) and the choice of resting background colors (behavior) were affected by the temperature experienced during development. We found that dry season temperatures lead to browner butterflies with a stronger preference to rest on brown backgrounds. While behavior is typically very flexible in relation to changes in the adult environment, a developmental ‘imprint’ on adult behavior can help ensure an effective match between the cryptic appearance and background choice behavior.

## 1. INTRODUCTION

Predation is an important selective pressure and organisms have evolved a wide range of adaptations to improve chances of survival in the presence of predators. Such adaptations typically include a suite of traits, including morphology and behavior, that work together to maximize predator avoidance. For example, spider-mimicking moths both look and act like their predators to improve their chances of deceiving them (Wang et al. 2017). Butterflies bearing large eyespots, generally only expose these when threatened to increase their chances of startling predators (Olofsson et al. 2012; De Bona et al. 2015). While defensive mechanisms are often constitutive properties of prey species (e.g. the spines of the hedgehog), they can also be induced by specific environmental conditions predictive of predation risk. For example, the protective morphology of *Daphnia* crustaceans is induced by chemical cues that signal predator presence (Grant and Bayly 1981; Weiss et al. 2015). Plasticity in anti-predator strategies also occurs in response to abiotic factors such as those that affect the visual background under which species need to escape predator attention. For example, in many polyphenic butterflies, seasonal strategies to avoid predation are associated with seasonal fluctuations in climatic cues which anticipate variation in background vegetation cover (Brakefield and Reitsma 1991). While the environmental regulation of development, or developmental plasticity, often leads to changes in adult morphology that are irreversible, adult behavior is generally very responsive to environmental conditions experienced during adulthood (Sih et al. 2004). The question of whether plastic responses of morphological and behavioral anti-predator traits are coordinated, to ensure that animals use a behavioral repertoire that matches their appearance, remains open.

The tropical butterfly *Bicyclus anynana* is a well suited experimental model to test hypotheses about the integration of multiple environmentally-induced anti-predator traits (Brakefield et al. 2009). This species exhibits polyphenism as an adaptive response to dry-wet seasonal environments, with crypsis and deflection as seasonally alternating anti-predator strategies (Brakefield and Reitsma 1991). Dry season individuals with small wing pattern elements are camouflaged against the brown background of dry leaves, while wet season individuals have large marginal eyespots that deflect predators’ attention away from the more vulnerable body (Beldade and Peralta 2017). This seasonal variation in wing patterns is induced by the temperature experienced during development, which anticipates seasonal changes in precipitation and vegetation coverage. The cooler temperatures of the dry season lead to the development of wings with smaller pattern elements (Kooi and Brakefield 1999). Mark-release-recapture experiments in the field (Brakefield and Frankino 2009) and experimental work with different putative predators (Lyytinen et al. 2004; Prudic et al. 2015) have provided support for the adaptive advantage of seasonal variation in eyespot size.

Here we use thermal plasticity in *B. anynana* and a full factorial design to study the combined effects of developmental and adult temperature on anti-predator pigmentation and behavior. While thermal plasticity for *B. anynana* eyespot size is well characterized, other traits that might enhance the crypsis of dry season individuals, such as actual wing color and background color preference, have largely been ignored (but see Brakefield and Reitsma 1991; Lyytinen et al. 2004). We investigated thermal plasticity in wing colors and in behaviors that are predicted to make crypsis more effective. These behaviors are: 1) choice for matching background colors, by which animals choose to rest on backgrounds that better resemble their appearance (Endler 1984; Michalis et al. 2017), and 2) motionlessness, since camouflage is known to be less effective when animals are in motion (Ioannou and Krause 2009). We test the hypotheses that the dry season form individuals, believed to be cryptic against the dry foliage of their season (Lyytinen et al. 2004; Brakefield and Frankino 2009), will not only have smaller pattern elements, but also colors that are less conspicuous. Conversely, highly contrasting ornaments in wet-season individuals could increase their effectiveness in deflecting predator attention away from the body (Kjernsmo et al. 2018). In addition, we hypothesize that the presumably more cryptic dry season individuals will be less active and will rest preferentially on brown backgrounds reminiscent of the natural dry foliage. Globally, this work explores the extent to which seasonal strategies for avoiding predation involve synchronized developmentally plastic responses across traits, to prevent a mismatch between anti-predatory appearance and behavior.

## 2. METHODS

### (a) Butterflies and rearing conditions

Eggs from a laboratory population of *B. anynana* (Brakefield et al. 2009) were allocated to one of two climate-controlled rooms (65% Relative Humidity, 12L:12D photoperiod) differing in ambient temperature to simulate the conditions of the dry (Td=19°C) and wet seasons (Td=27°C). Groups of 250 larvae were reared in large population cages and fed with young maize plants. Upon eclosion, males and females were separated and allocated to one of two adult temperature regimes (Ta; 19°C or 27°C). This resulted in four treatments with different combinations of developmental (Td) and adult (Ta) temperatures (Fig. S1). Adults were kept at densities of 25 same-sex individuals per cage, and fed with moist banana.

### (b) Wing pigmentation analysis

Using a color-calibrated digital scanner (Epson V600) that provides a linear response to changes in light levels (Stevens et al. 2007; Fig. S2), we scanned the ventral surfaces of hindwings of 16 female and 16 male butterflies per treatment (N=128 individuals). The resulting images were analyzed with custom-made interactive Mathematica notebooks (Rodrigues et al. 2017). We first drew two contiguous transects defined by five landmarks on the wing compartment bearing the eyespot typically used to characterize seasonal polyphenism in *Bicyclus* and related genera (van Bergen et al. 2017). We then specified the limits of the central band and each of the eyespot color rings along the transect. The colors of the wing background and different pattern elements were quantified by extracting the mean RGB values of 3×3 pixel squares centered on the transect (Fig. 1*a*). Data were then converted to CIE-xyY coordinates and plotted in a CIE-xy chromaticity diagram, which is a normalized representation of the colors perceived by trichromatic observers (Stevens et al. 2009; Fig. S3). For each individual, we calculated the Euclidean distance between the color of the different pattern element components and that of the background of the wing. These distances were used as a proxy for internal contrast, or conspicuousness of pattern element components in relation to the wing background. We also calculated the distance from each pixel along the transect to the brown patches used in our behavioral assay (details below). The mean distances were used as an inverse proxy for color similarity or crypsis in our experimental setup.

**Figure 1.**
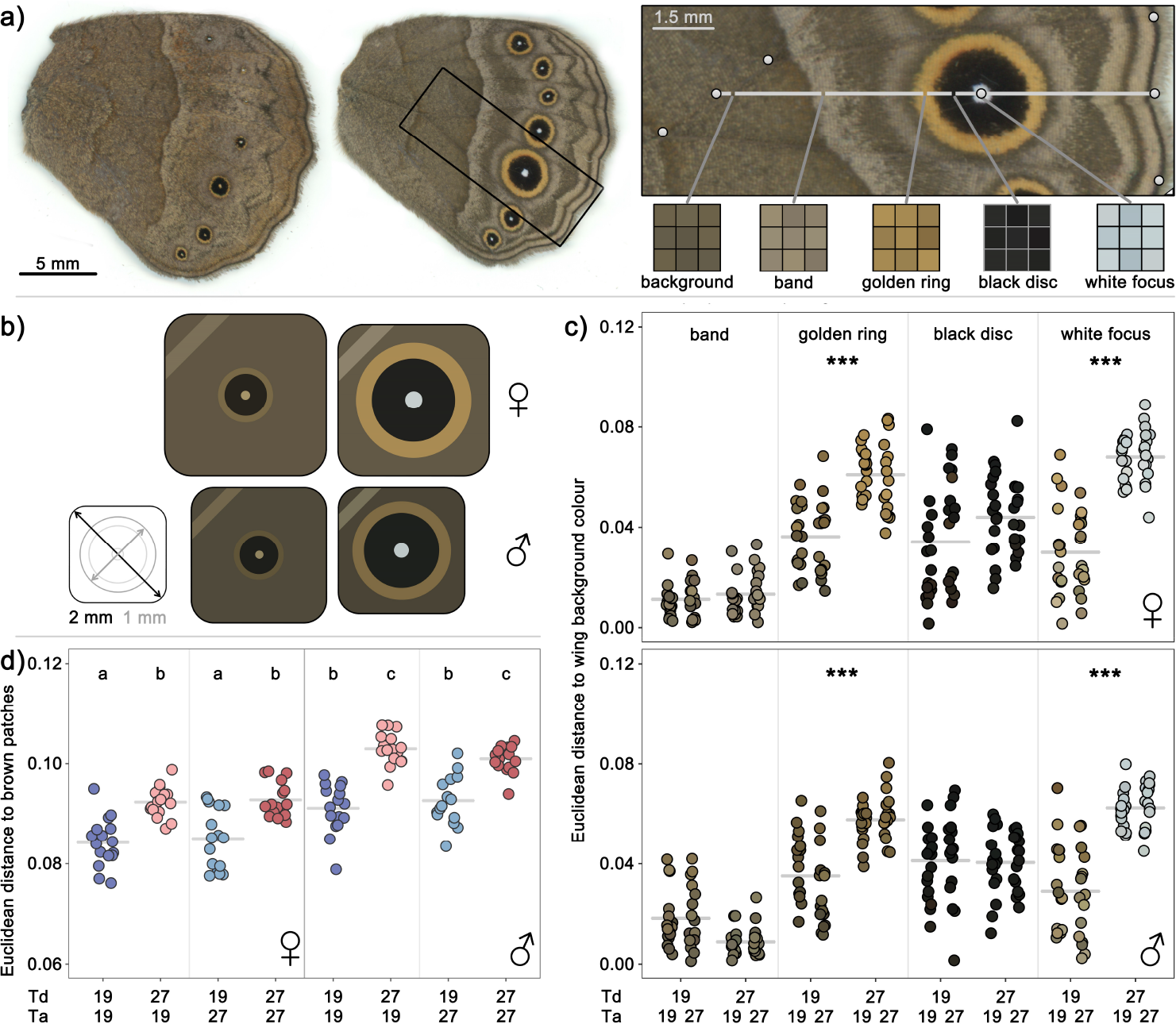
Environmentally-dependent wing pigmentation. (*a*) Examples of ventral hindwing surfaces from females reared at Td=19°C (left) and Td=27°C (right; with section highlighted). The section of a hindwing shows landmarks (white circles) that defined two contiguous transects (white line) passing through the center of the eyespot. The colors of the wing background, central band, and eyespot rings were quantified by extracting the mean RGB values of 3×3 pixel squares. Data were converted to CIE-xyY color space prior to color analyses. (*b*) Schematic representation of the color of the background (square fill), central band (stripe on top left corner) and eyespot rings (circles), as well as the size of the wings and eyespot rings (cf. scale on left-most drawing), for individuals reared at Td=19°C (left) and Td=27°C (right). (*c*) Euclidian distance in a CIE-xy chromaticity diagram between the color of the background and the color of different pattern elements for females and males from four thermal treatments. Color contrast was not affected by adult temperature (Ta), statistical significance for effect of developmental temperature (Td) is indicated as *** for P<0.001. (*d*) Mean Euclidian distance in a CIE-xy chromaticity diagram, using all pixels along the transect, to the color of the brown patches used in our behavioral assay. Significant differences between groups (Tukey’s HSD, P<0.05) are indicated by different letters. Details of statistical tests in Table S1.

### (c) Behavioral assays

The behavioral assays were performed in large flight cages (150×50×54 cm) that allow expression of a more natural behavior of experimental *B. anynana* (Joron and Brakefield 2003). The bottoms, backs, and sides of those cages were covered with canvas depicting a mosaic of green (Hue=130°, Saturation=60%, Brightness=50%) and brown (HSB 25°, 60%, 50%) squares (16.7 cm^2^). The fronts were covered with mesh netting and the tops had three fluorescent lamps (Osram L58W/840). Digital cameras (Canon EOD 1000D), connected to a Wireless Remote Control (Hähnel - Giga T Pro), were positioned in front of the cage. Cohorts of 20 same-sex, 6-8 day old individuals were released into the flight cage and left to habituate for two hours. Subsequently, about 8 hours after the lights were switched on, 15 time-lapse photos were taken at four-minute intervals for a period of 60 minutes. For each photo, we registered the number of butterflies at rest on each of the two background colors, as well as the number of individuals resting at exactly the same position as in the previous time point (i.e. no re-location). For perching preference, we used the number of new arrivals to brown patches and the number of new arrivals to green patches, for each time point, as a 2-vector dependent variable. This binomial variable was interpreted as a preference for brown patches, but could, conversely, also be seen as a tendency to avoid green backgrounds. For activity levels we used the number of inactive individuals and the number relocated individuals as a 2-vector dependent variable. Sixteen cohorts were tested for each experimental treatment, eight for each of the sexes (N=64 cohorts).

### (d) Statistical analyses

All statistical tests were done using R (R Core Team 2017) and the package *afex* (Singmann et al. 2018). We examined the effects of developmental temperature (Td), adult temperature (Ta), and sex on wing pigmentation using linear mixed models (Table S1), and on perching preference and activity levels using generalized linear mixed models with a binomial distribution and a logit link. For the behavioral data, the cohort and the time point (i.e. consecutive photos) were included as random effects in the models. For both behavioral variables, minimum adequate models were obtained by backward elimination, starting from the full model and using the Bayesian Information Criterion (BIC). P-values were calculated via likelihood ratio tests and by parametric bootstrapping (nsim=999). Post-hoc pairwise comparisons (Tukey’s HSD; alpha=0.05) were performed using the package *lsmeans* (Tables S2 and S3).

## 3. RESULTS

We used a full factorial design, with two developmental temperatures (Td; 19°C and 27°C) and two adult temperatures (Ta; 19°C and 27°C) that simulate the natural dry and wet seasons, respectively. This enabled us to study thermal plasticity in pigmentation and behavior in the seasonal plastic butterfly *B. anynana*, and to partition environmentally-induced variation into early and late life effects.

### (a) Thermal plasticity in wing colors

We quantified the color of different wing pattern elements (central band and eyespot rings; Fig. 1*a*), as well as of the wing background of adults from each of the four combinations of Td and Ta. We found that developmental temperature affected many of these aspects of wing pigmentation in both males and females. Pattern elements of individuals reared under cooler temperatures were not only smaller, as had been shown previously (Kooi and Brakefield 1999; Mateus et al. 2014), but they were also of less contrasting colors. Specifically, the colors of the eyespot’s external ring (Td; F=114.23, P<0.001) and central focus (Td; F=186.29, P<0.001) were significantly closer to the background color of their respective wings (i.e. eyespot colors were ‘browner’; Fig. 1*b,c*) in individuals reared at 19°C relative to those reared at 27°C. Moreover, overall wing coloration of individuals reared at 19°C was closer to the brown background patches used in our behavioral assays (Td; F=134.70, P<0.001), with the colors of females being lighter and closer to the brown patches relative to those of males (Sex; F=115.02, P<0.001; Fig. 1*d*). Adult temperature had, as expected, no effect on wing pigmentation (Table S1).

### (b) Thermal plasticity in color preference

The behavior of males and females from the four temperature treatments was monitored in large flying cages containing a mosaic of green and brown patches. We monitored their preference to rest on brown versus green (i.e. perching preference), as well as the likelihood of re-locating between sequential time points (i.e. levels of activity) (Fig. S4). We found that developmental temperature affected adult color preference (Fig. 2*a*). Compared to their wet-season counterparts (Td=27°C) with more contrasting wing color patterns, dry-season individuals (Td=19°C) were more likely to alight in brown patches (Td; χ^2^=20.66, P<0.001). This was true for both sexes and in both adult temperatures. The preference for brown backgrounds, or the avoidance of green ones, was stronger in females than in males (Sex; χ^2^=62.53, P<0.001), and these sex-specific color preferences were stronger when animals were tested at lower adult temperatures (Ta:Sex; χ^2^=14.26, P<0.001). Activity levels of females were not affected by either Td or Ta while, in contrast to our expectation, males were more active when reared (Td:Sex; χ^2^=7.79, P=0.005) or kept (Ta:Sex; χ^2^=22.91, P<0.001) at cooler temperatures (Fig. 2*b*).

**Figure 2.**
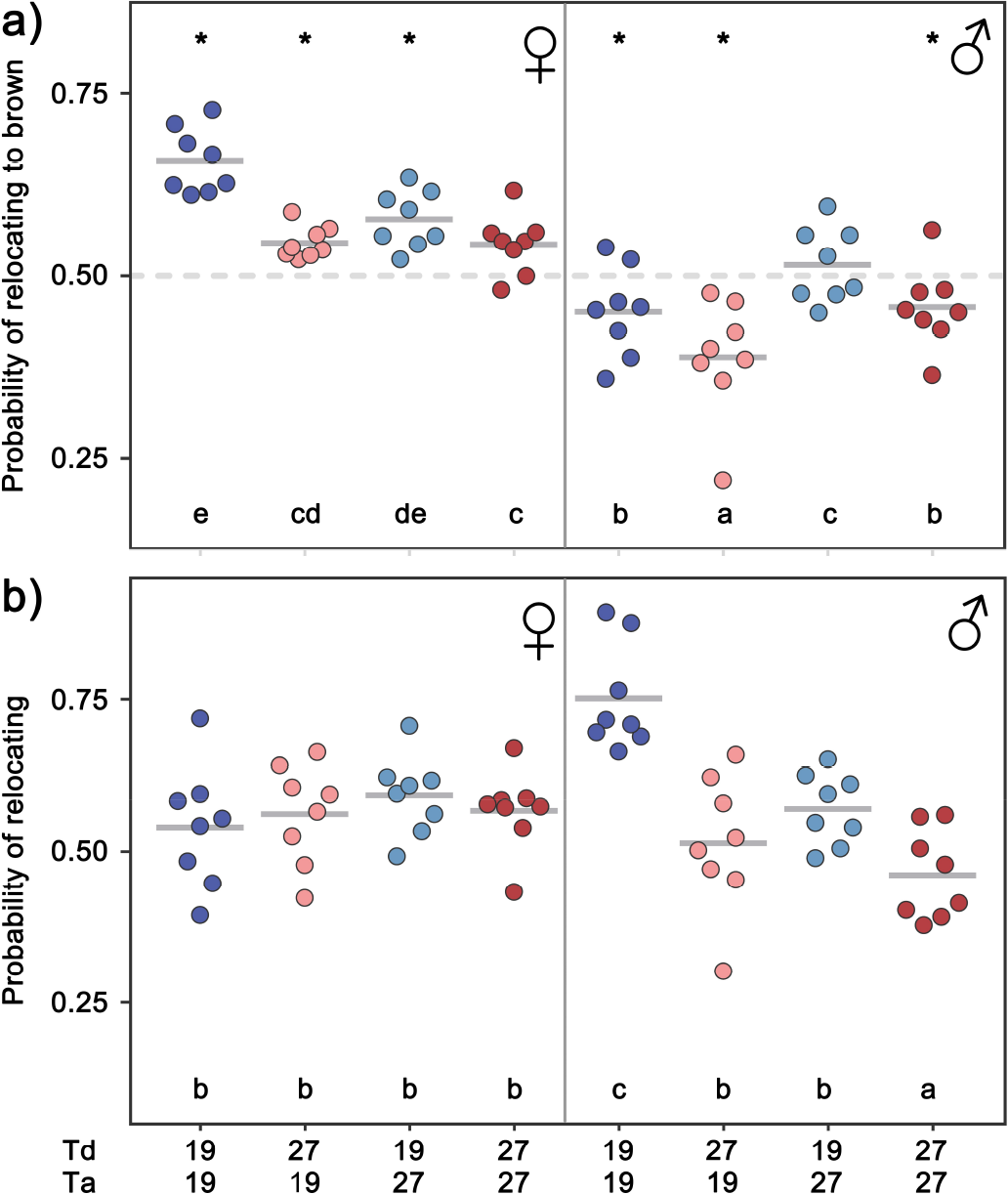
Environmentally-dependent behaviors. (*a*) Probability of relocating to a brown patch for females and males from different combinations of developmental and adult temperature (Td and Ta, respectively). Within the same testing temperature and sex, dry season individuals (Td=19°C; blue dots) are more likely to alight on brown patches than individuals with a more conspicuous coloration (Td=27°C; red dots). Preference for brown patches is stronger in females and sex-specific color preferences are stronger when individuals were tested at cooler temperatures (Ta=19°C). Asterisks indicate groups having probabilities of relocating to brown that are significantly different from a 1:1 ratio. (*b*) Probability of relocating for females and males from different treatments. Males were more active when reared or kept at cooler temperatures while activity levels of females were not affected by either Td or Ta. Dots in both panels represent the probability of relocating (to brown) for each cohort based on the total number of relocations (to brown) observed throughout the assay. Significant differences between groups (Tukey’s HSD, P<0.05) are indicated by different letters. Details of statistical tests in Tables S2 and S3.

## 4. DISCUSSION

Using a full factorial design that allows separating the effects of developmental and adult environments (Groothuis and Taborsky 2015), we showed that both adult appearance and adult behavior in *B. anynana* are affected by the temperature experienced during development. We demonstrated that developmental temperatures affect wing pigmentation beyond the size of the ornamental pattern elements. In accordance with dry season individuals relying on camouflage against the dry brown foliage to avoid attracting predator attention, they not only develop smaller pattern elements (eyespot and central band, as previously documented), but the colors of these elements contrast less with their wings’ brown background color (Fig. 1). Conversely, this also matches the expectation that selection for a deflective effect of marginal eyespots in the wet season should favor the more contrasting eyespot colors we documented for wet-season individuals (see also Kjernsmo et al. 2018). Moreover, we also showed that individuals reared at cooler temperatures (i.e. the browner ones) also had stronger preferences for brown resting backgrounds (Fig. 2*a*). Such a concerted response in developmental effects on adult color and color preference can help ensure that the development of a cryptic adult appearance is matched by a behavioral repertoire that enhances crypsis (for a review, see Stevens and Ruxton 2018).

The perching preference plasticity that we demonstrate here may reduce the risk of detection by predators when cryptic individuals are confronted with visually heterogeneous environments (Merilaita et al. 2001). For *B. anynana* this is especially relevant during seasonal transitions when dead brown leaves and green vegetation co-occur, and individuals can choose among a variety of backgrounds to rest on. We also found that the preference for brown backgrounds was stronger in females. In general, regardless of developmental (Td) or adult (Ta) temperatures, males were more likely to perch on green patches (Fig. 2*a*). This sexually dimorphic preference for resting backgrounds may reflect that males and females adopt different habitat use strategies in the wild, as documented for other species (eg. Shine 1986; Merilaita and Jormalainen 1997). Males of *B. anynana* pursue a perch-and-chase strategy to procure mates and when conditions are favorable for reproduction they are typically found in patches of green sunlit vegetation that improve the likelihood of detecting potential mates (Vande Velde et al. 2011; van Bergen et al. 2013). In contrast, females tend to fly more closely to the ground, which is necessarily browner, searching for suitable host plants for oviposition which are available from the start of the wet season onward (Brakefield and Reitsma 1991). While our data support the hypothesis that more uniformly brown individuals show stronger preference for the background color that presumably maximizes camouflage (i.e. browner butterflies tend to perch on brown resting sites), we found no evidence for responses in another type of behavior that could potentially enhance crypsis; motionlessness. Field observations had suggested that cryptic adults spent the dry season being relatively inactive, probably to conserve energy and avoid detection (Brakefield and Reitsma 1991). However, we found no evidence for reduced activity in dry-season-like individuals in this study. Though we aimed to use experimental conditions that allow for the expression of more natural behaviors, it is unclear to what extent the behavioral traits quantified here could be affected by factors such as the presence of (same-sex) con-specifics, the absence of real sunlight, and attributes of the color patches used (e.g. their sizes, shape, and actual colors).

We demonstrated that environmental conditions during early life can affect wing coloration and can also lead to “fixed” changes in adult behavior, as had been shown in insects and in vertebrates (e.g. Bear et al. 2017; Fischer et al. 2017). While the mechanisms underlying the environmentally-induced differences in *B. anynana* color and color preference were not explored here, it is likely that traits responding in an integrated manner to environmental cues share underlying regulators. Traits making up the *B. anynana* seasonal syndrome, including eyespot size (Mateus et al. 2014), life-history traits (Oostra et al. 2014), and courtship behavior (Bear et al. 2017), are known to be affected by developmental temperature-dependent ecdysteroid hormone dynamics during the late-larval and early pupal stages. How ecdysone dynamics can affect the biosynthetic pathways for pigment production in *B. anynana* (Beldade and Peralta 2017; Zhang et al. 2017) is unclear, but effects of developmental temperature on these pathways have been studied in detail in *Drosophila melanogaster* (Gibert et al. 2007, 2017; De Castro et al. 2018), a well-described model for thermal plasticity in body pigmentation. Studies in other systems have also shown how temperature-induced changes in ecdysteroid signaling can affect adult behavior in different manners: by altering neuronal activity and connectivity (e.g. in *D. melanogaster*; Ishimoto et al. 2013; Carvalho and Mirth 2015) and/or by mediating visual sensitivity of adults through the regulation of eye development (e.g. in *Manduca sexta*; Champlin and Truman 1998). In fact, the size of the *B. anynana* compound eyes is known to be both developmentally plastic and sexually dimorphic (Everett et al. 2012). Wet-season individuals and males have larger eyes relative to dry-season individuals and females, respectively. As variation in eye morphology can impact visual performance (e.g. Land 1997; Merry et al. 2006) we speculated that seasonal differences in eye size could affect color vision and color preferences. We investigated the relationship between thermal plasticity in eye size (published data in Everett et al. 2012) and our own preference behavioral data. Curiously, we found that individuals reared at lower temperatures have both smaller eyes and stronger preferences for brown resting backgrounds (Fig. S5).

Overall, our data clearly show that pigmentation and behavioral anti-predator traits in *B. anynana* are affected in concert by the conditions experienced during development. The coordinated responses to developmental temperature are consistent with favoring a match between appearance and behavior that maximizes the effectiveness of crypsis as an alternative seasonal strategies for predator avoidance. Globally, our study underscores the importance of a tight integration of form and function necessary for survival under heterogeneous environments. The coordination of morphology and behavior is of prime importance and evolutionary changes in each type of trait — shaped by multiple factors, including predator avoidance — can potentially promote change in the other.

## ACKNOWLEDGEMENTS

We are grateful to Carolina Peralta for technical assistance, Filipa Alves for developing a set of interactive *Mathematica* notebooks we based our pigmentation phenotyping scripts on, Martin Stevens for valuable advice on color quantification, and two anonymous reviewers for suggestions which greatly improved our manuscript. Financial support was provided by the Portuguese science funding agency, *Fundação para a Ciência e Tecnologia*, FCT (PTDC/BIA-EVF/0017/2014); French research funding agency, *Agence Nationale de la Recherche*, ANR (Laboratory of Excellence TULIP, ANR-10-LABX-41) and French research center, *Centre National de la Recherche Scientifique*, CNRS (International Associated Laboratory, LIA BEEG-B). The authors declare no conflicts of interest.

## AUTHORS’ CONTRIBUTIONS

EvB and PB conceived and designed the experiments. EvB performed the experiments and analyzed the data. EvB and PB wrote the manuscript.

## DATA ACCESSIBILITY

Raw data will be archived in the Dryad Digital Repository upon acceptance of the manuscript.

## SUPPLEMENTARY MATERIALS

**Supplementary Figure 1.**
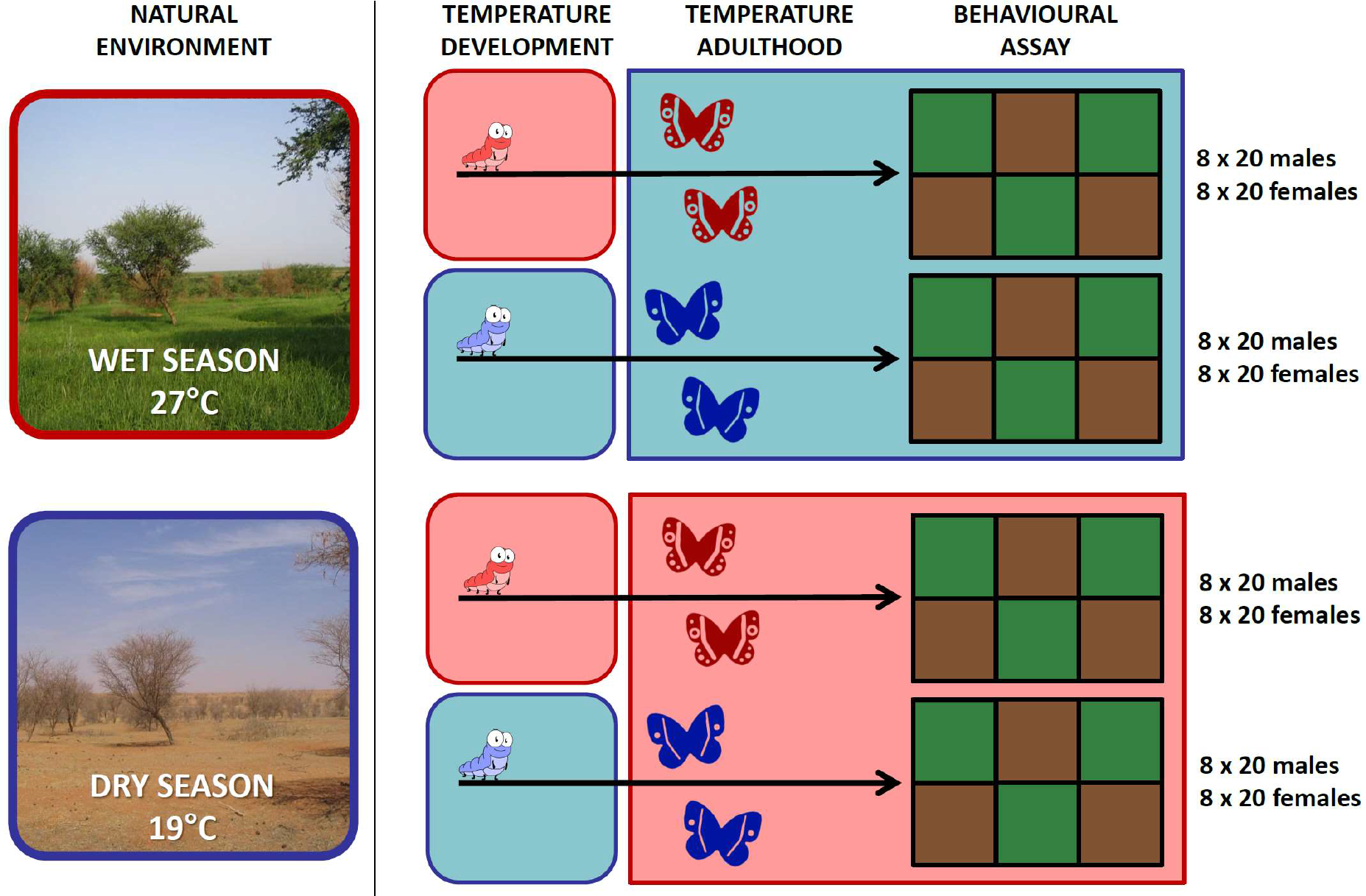
Schematic representation of the experimental design. Panels on the left-hand side depict the natural environment of *B. anynana* during the wet season (upper) and dry season (lower) while panels on the right-hand side provide a schematic representation of our experimental design. Individuals were reared, from egg to adult eclosion, at one of two developmental temperatures (Td; 19°C and 27°C) that simulate the ambient temperatures of the dry and wet season, respectively. Variation in rearing temperature is sufficient to induce seasonal polymorphism in this species, with cool and warm temperatures inducing dry season form (DS) and wet season form (WS) individuals, respectively. On the day of eclosion, the butterflies were allocated to one of two adult temperatures (Ta; 19°C and 27°C), yielding four experimental treatments with different combinations of Td and Ta. Behavioural assays were performed at the temperature of the adult regime using cohorts of 20 six to eight day old males or females (N_cohort_= 64). The red colours in this scheme represent warm temperatures or individuals reared at warm temperatures. Cool temperatures, or individuals reared at cool temperatures, are represented by blue colours. Photographs of natural environments are courtesy of Jonas Ardö.

**Supplementary Figure 2.**
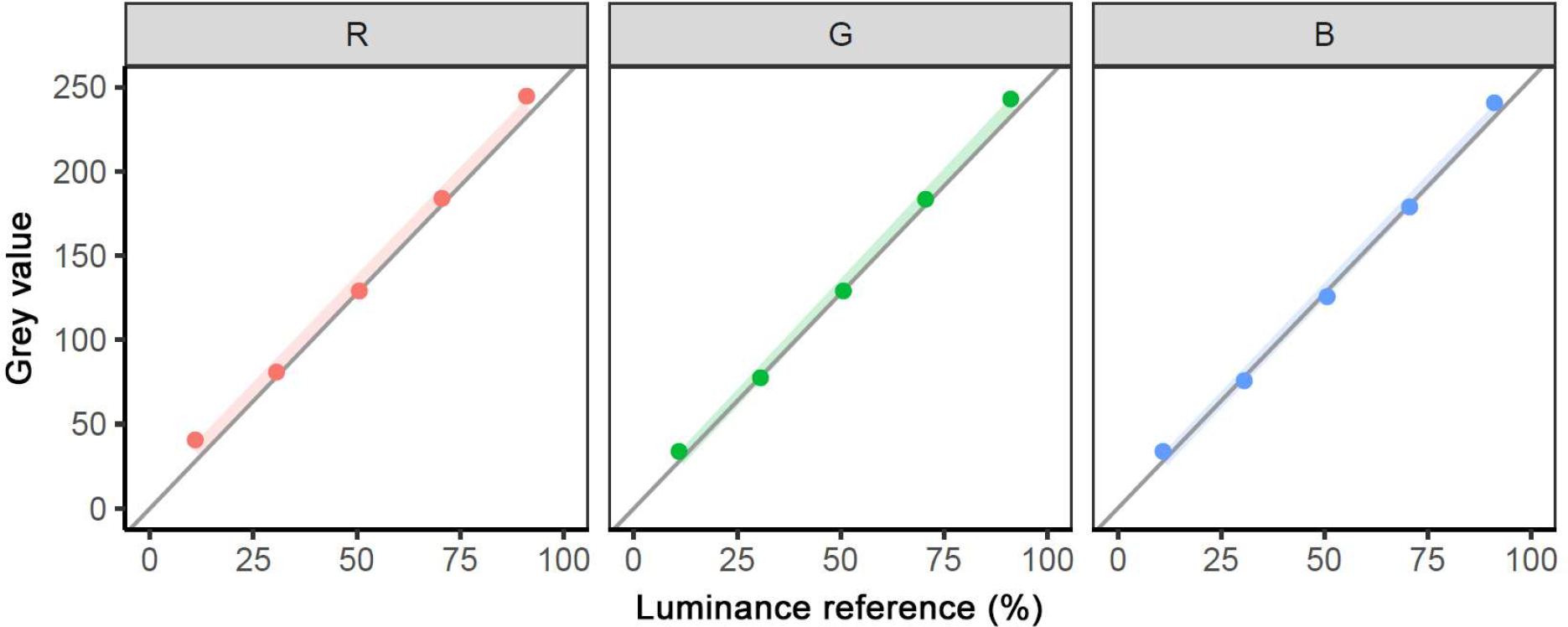
Colour calibration and linearisation of the scanner. The digital scanner (Epson V600) used to acquire wing colour information was calibrated using a colour reference card (Wolf Faust, R110714). The relationship between the grey scale value measured for a set of five reflectance standards (GS0, GS5, GS10, GS15 and G20 on the colour reference card) shows that the gamma curves are linear with a close fit to the required values (grey lines) in each of the three colour channels; R (longwave or ‘red’), G (mediumwave or ‘green’) and B (shortwave or ‘blue’).

**Supplementary Figure 3.**
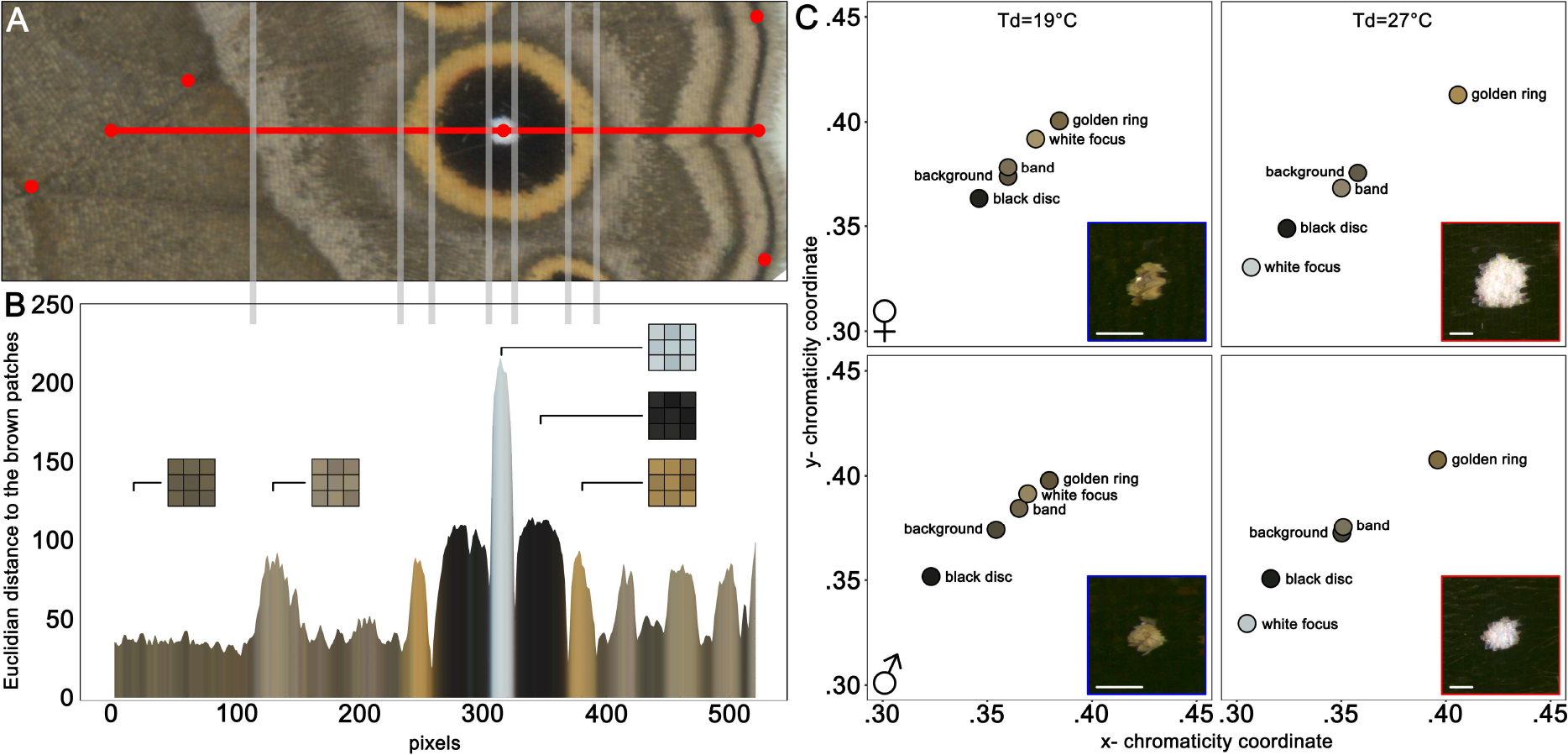
Quantification of crypsis-enhancing pigmentation traits. **A.** We used a set of custom-made interactive Mathematica notebooks to analyse the ventral hindwing surfaces of 16 female and 16 male butterflies per experimental treatment (i.e. two individuals per cohort, N=128). Two contiguous transects were defined by the centre of the large fifth eyespot (cell Cu1) and four wing landmarks (on the wing margin and intersection between veins). The four wing landmarks were used to calculate midpoint values along the wing vein and margin, and these midpoints determined the start and end of the contiguous transects. Secondary landmarks, represented by grey lines, were used to specify the edge of the central band and the limits of each of the colour rings along the transect line. **B.** The colours of the eyespot rings were quantified by extracting the mean RGB values of 3×3 pixel squares centred on the intersection between the midpoints of the secondary landmarks and the transect line. The mean RGB values of the background and central band were extracted from a 3×3 cluster located 10 pixels after the associated secondary landmarks on the marginal side. Here, each pixel along the transect is represented as its Euclidean distance, in RGB colour space, to the colour of the brown patches used in our behavioural assays. **C.** RGB values were converted (to CIE-xyY colour space) and represented in CIE-xy chromaticity diagrams (a normalized representation of the colours perceived by trichromatic observers). To estimate how similar or dissimilar the colours of wing pattern elements are we calculated the Euclidean distance between the colour of the each pattern element and the colour of the wing’s background. CIE-xy chromaticity diagrams give the mean values for the each pattern element for females (upper) and males (lower) reared at cool (Td=19°C; left) and warm (Td=27°C; right) temperatures. The inset in each plot represent a detailed image of the wing scales that make up the white focus of the eyespot.

**Supplementary Figure 4.**
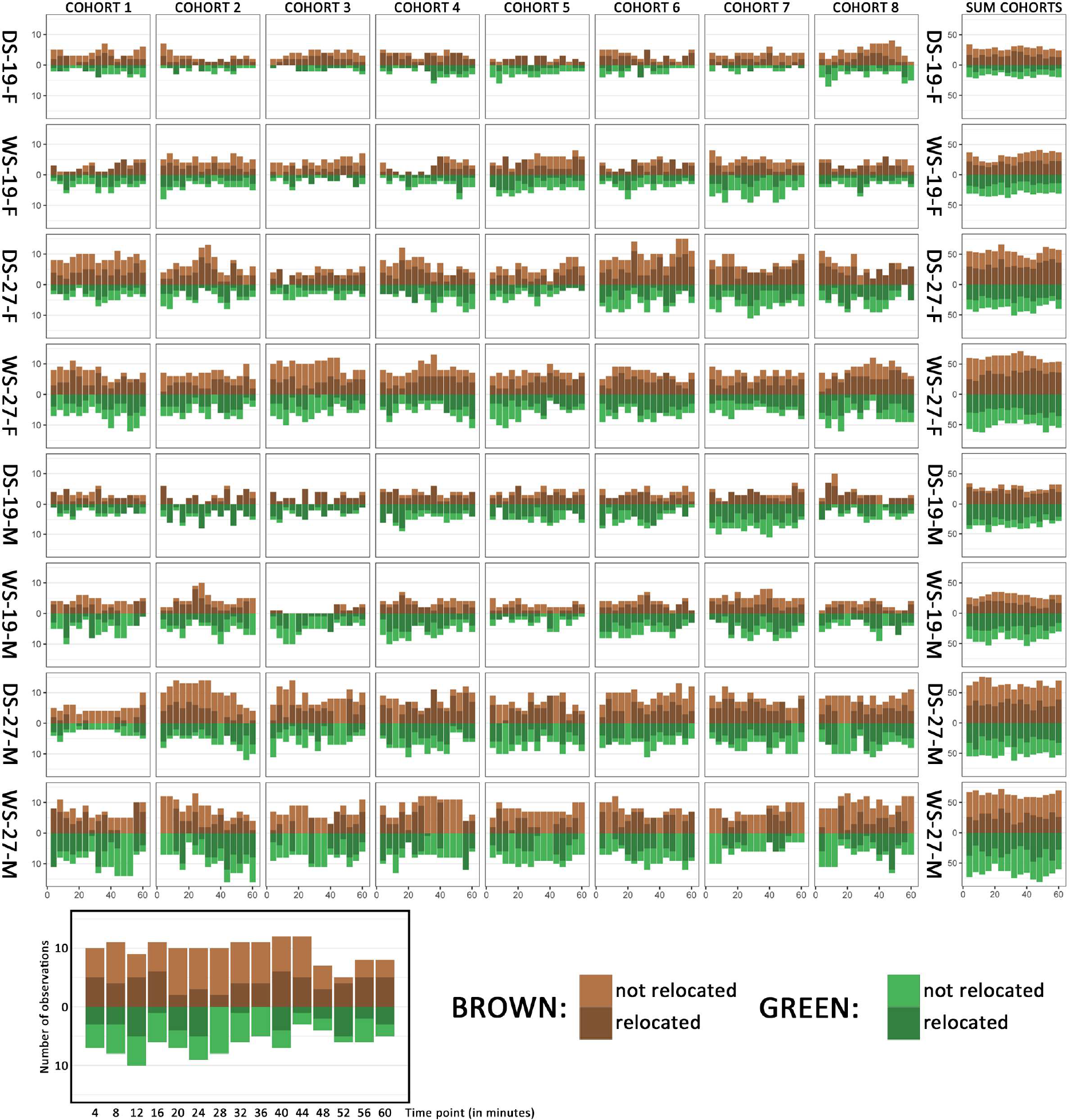
Behavioural data per cohort. Panels represent all behavioural data per cohort. The number of butterflies at rest on each of the two background colours is represented by the brown and green bars for each time point. The number of individuals resting at exactly the same position as in the previous time point (i.e. no re-location) is depicted by lighter colours inside each bar. Note that individuals that were resting on the mesh netting that covered the front of the cage or on the upper surface of the cage, or were airborne when the time-lapse photos were taken were denoted as ‘unaccounted for’ in the data. The number of individuals that was unaccounted for was higher at cooler temperatures (Ta=19°C) for both males and females.

**Supplementary Figure 5.**
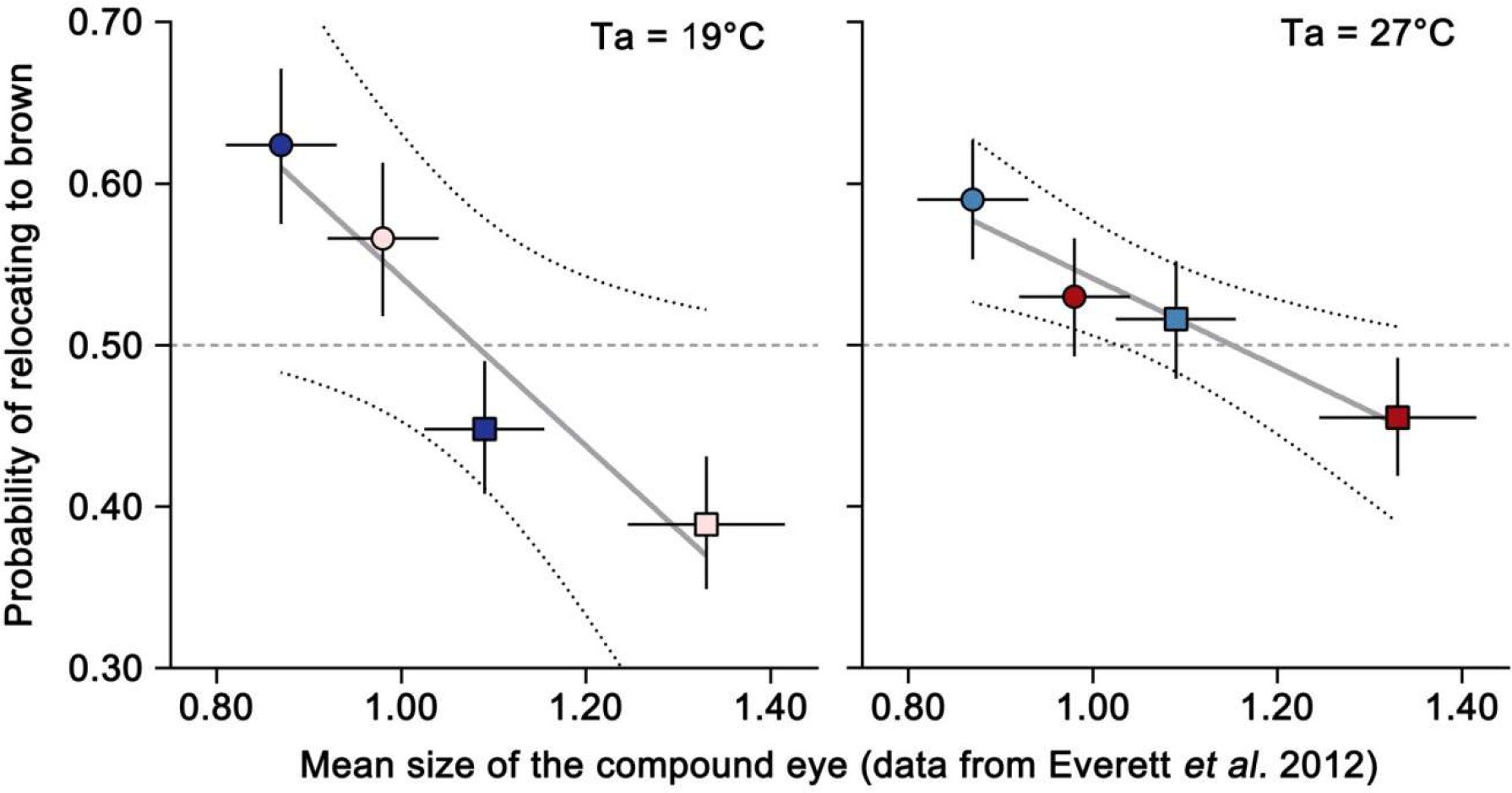
Correlations between eye size and perching preference. The size of the *B. anynana* compound eye is known to be developmentally plastic and sexually dimorphic. Individuals reared at warm temperatures (wet-season; red dots) have larger eyes than individuals reared at cold temperatures (dry-season; blue dots) and females (circles) have smaller eyes than males (squares); data from Everett *et al.* 2012. Our behavioural data on perching preferences (y-axis) correlates strongly with the published data on thermal plasticity in eye size (x-axis) such that individuals with smaller eyes have stronger preferences for brown patches. Left-hand panel gives the correlation between eye size and perching preferences measured at cool temperatures (Ta=19°C; *r*=−0.522, *R*^2^=0.914, P=0.044) while the right-hand panel shows the correlation with behavioural responses assessed at warm temperatures (Ta=27°C; *r*=−0.274, *R*^2^=0.949, P=0.026).

**Supplementary Table 1.**
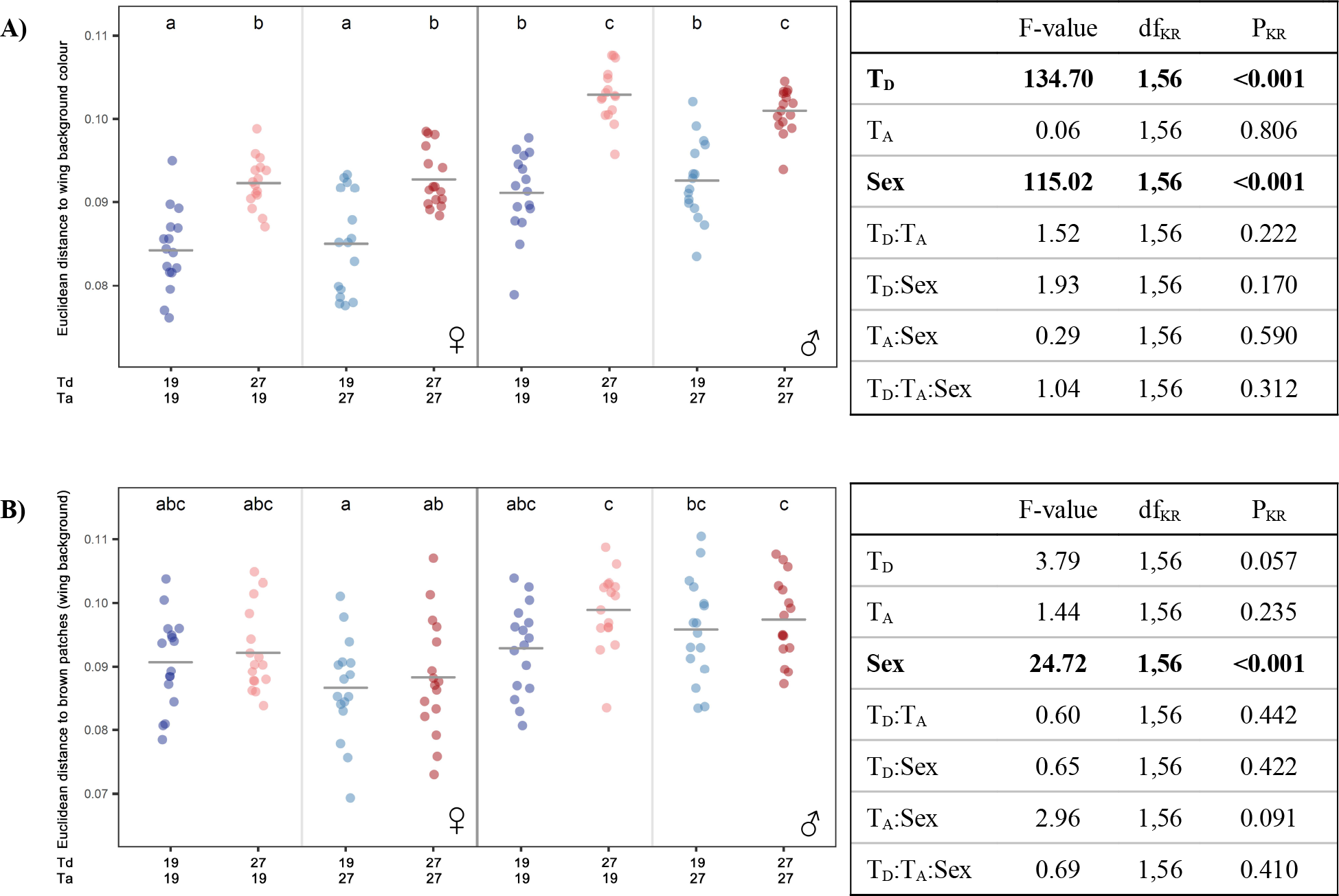
Colours and sizes of the wing pattern elements across treatments. Left-hand panels represent the Euclidian distance in CIE-xyY colour space from the colour of the brown patches used in the experiment to **A)** all pixels along the transect and **B)** the background colour of the wing. Panels C to F represent the Euclidian distance from the background colour of each individual to **C)** the central band, **D)** the golden ring, **E)** the black disc and **F)** the white focus. Panels G to K are related to (relative) sizes and panels represent **G)** the length of the transect (i.e. proxy for wing size), **H)** the relative distance from the start of the transect to the proximal edge of the central band (i.e. inverse proxy for the width of the band, which is difficult to measure directly because of its indistinct distal edge), **I)** the relative width of the golden ring, **J)** the relative width of the black disc and **K)** the relative width of the white focus. Relative widths were calculated by dividing the absolute size by the length of the transect. Tables on the right-hand side of each panel give the effects of developmental temperature (Td), adult temperature (Ta), sex and their interactions on the phenotypic trait measured. Effects with P<0.01 are shown in bold. The experimental cohort was included as a random factor and denominator degrees of freedom were determined with the Kenward-Roger method, using the package *afex*. Post-hoc pairwise comparisons (alpha=0.05) were conducted using the package *lsmeans*. Differences between experimental treatments and sexes are given in the graphs, with groups that are not significantly different from each other sharing a common letter.

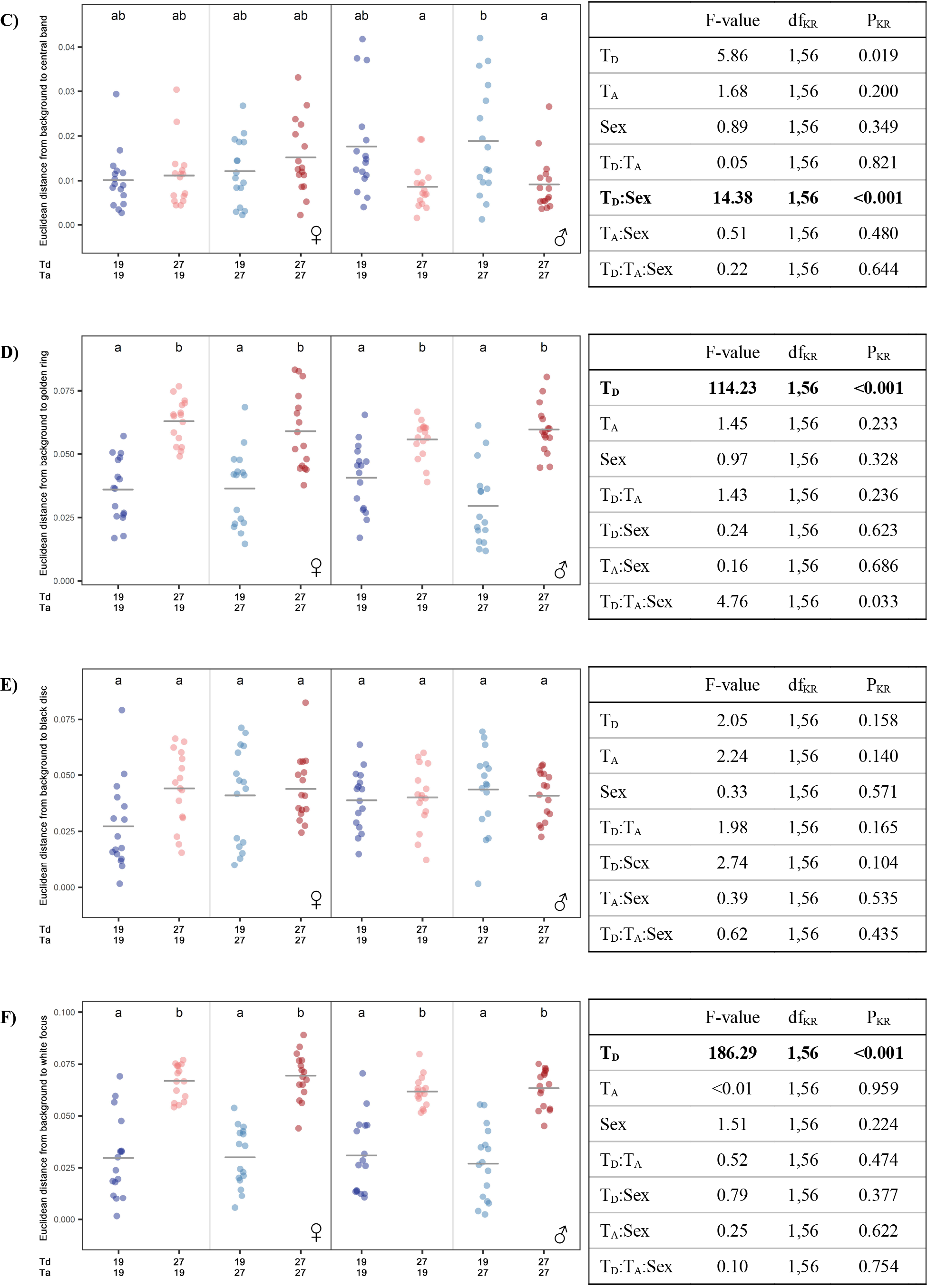

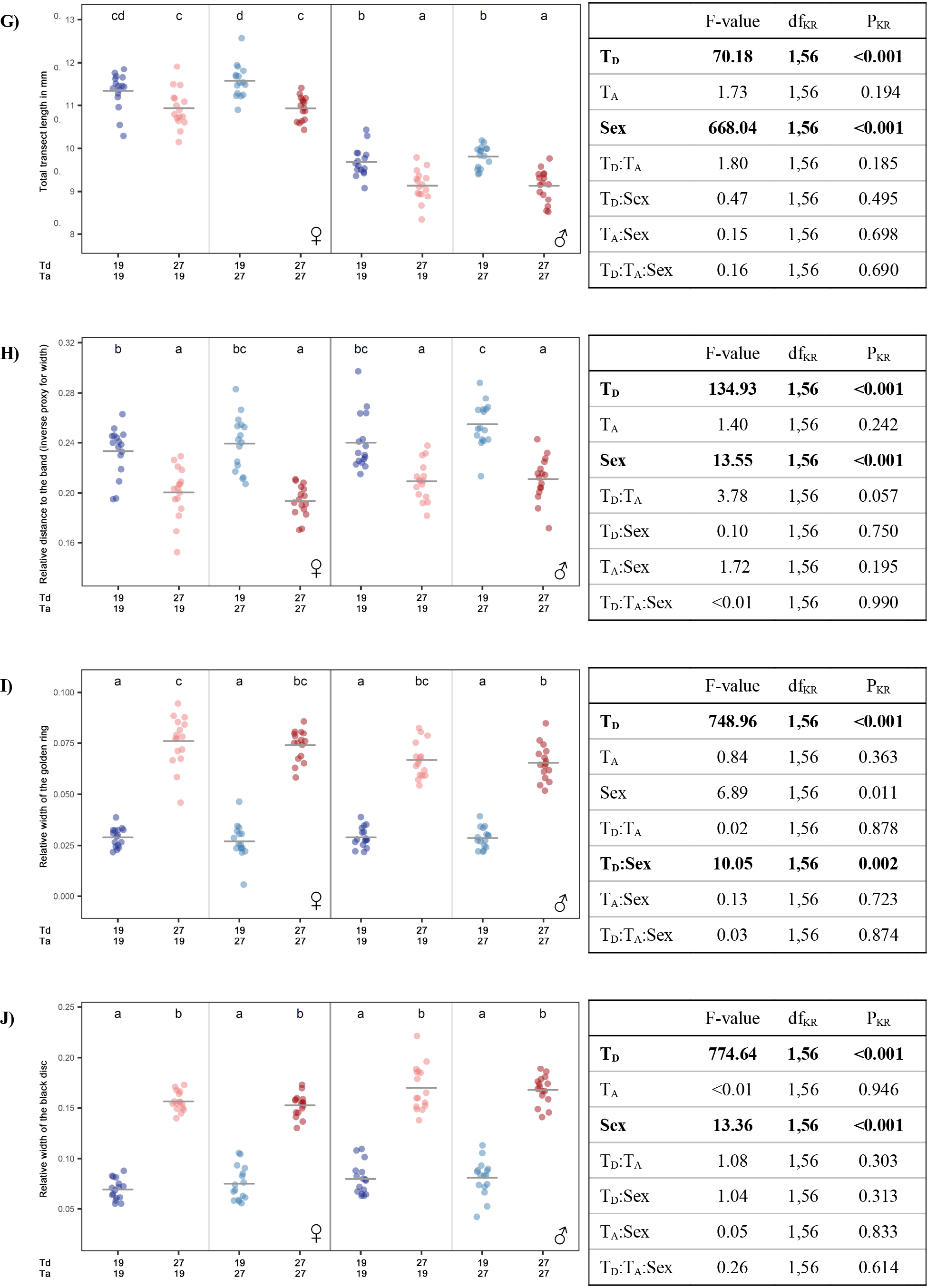

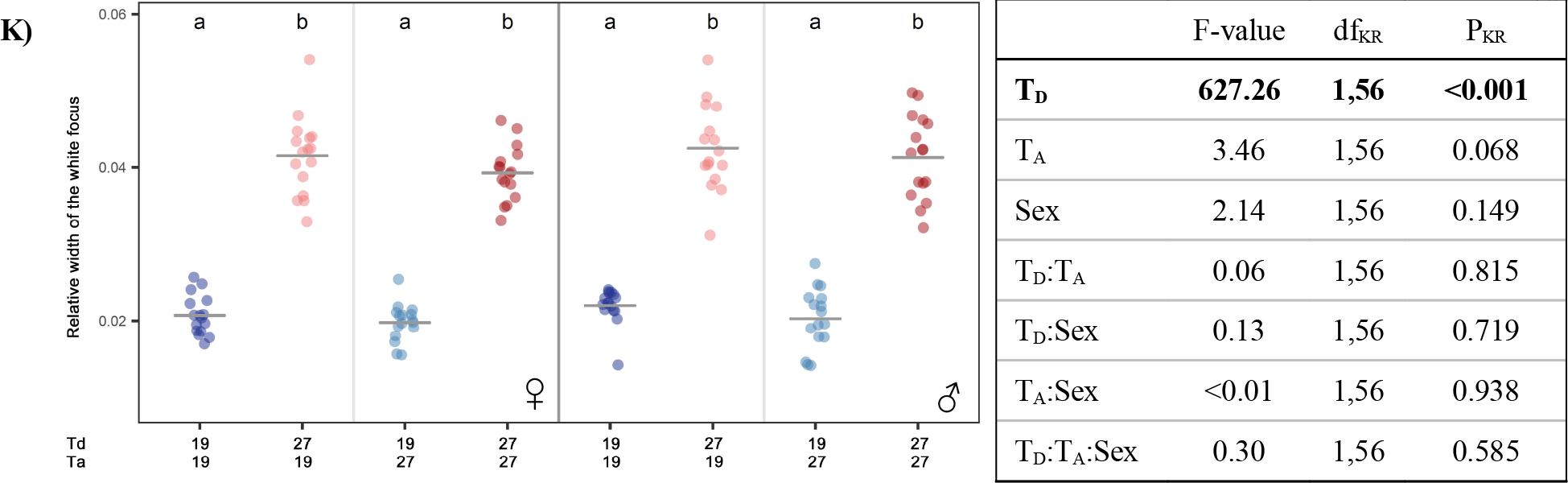

**Supplementary Table 2:**
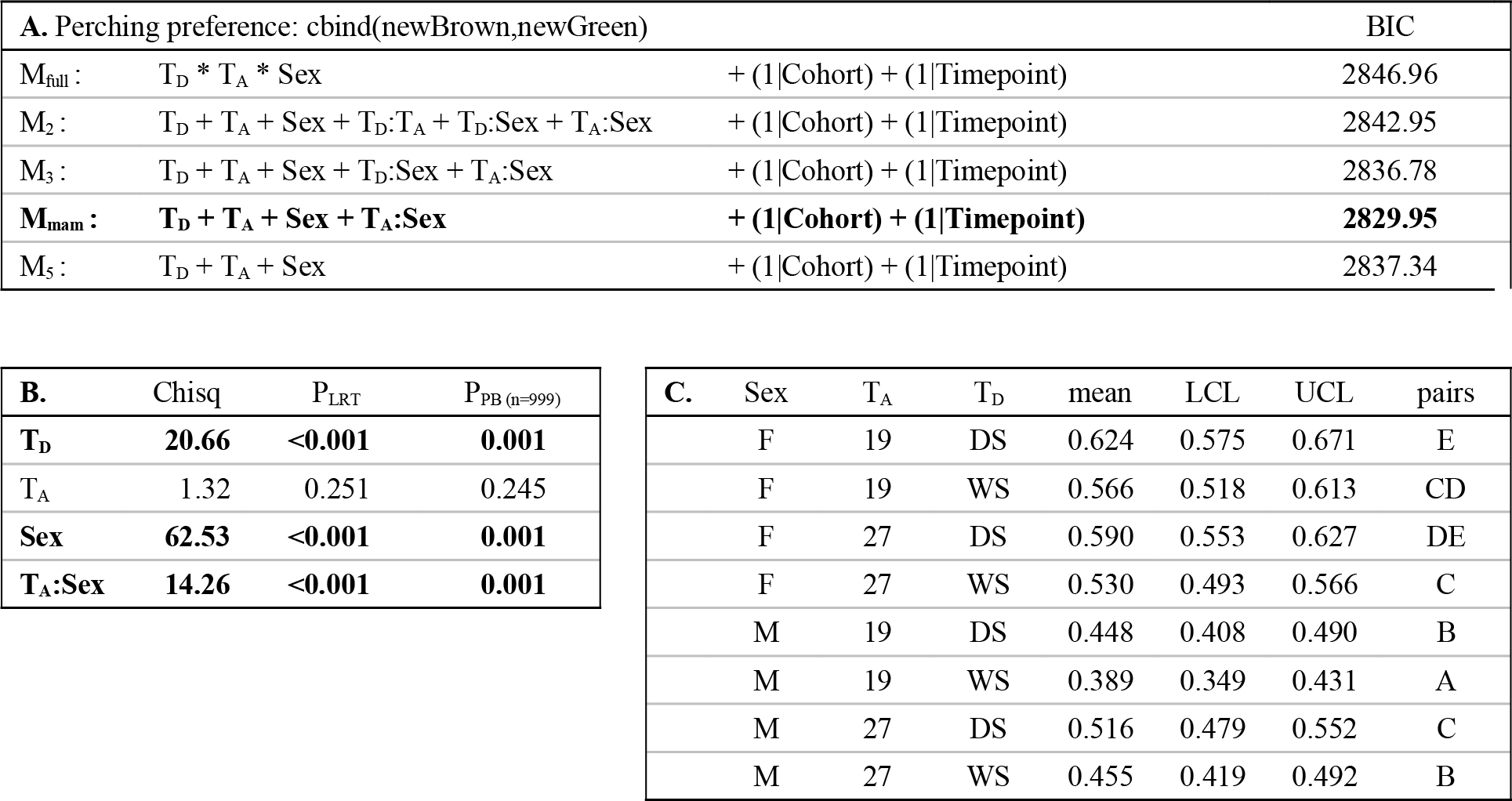
GLMM for perching preference (related to Fig 2a in the main text). **A.** Minimum adequate models were obtained by backward elimination, starting from the full model and using Bayesian Information Criterion (BIC). **B.** P-values for the minimum adequate model were calculated via likelihood ratio tests and parametric bootstrap (nsim=999), using the package *afex*. **C.** Mean probabilities, as well as upper and lower confidence limits, are given for each sex and experimental treatment. Post-hoc pairwise comparisons (alpha=0.05), obtained using the package *lsmeans*, specify differences between experimental treatments and sexes.

**Supplementary Table 3:**
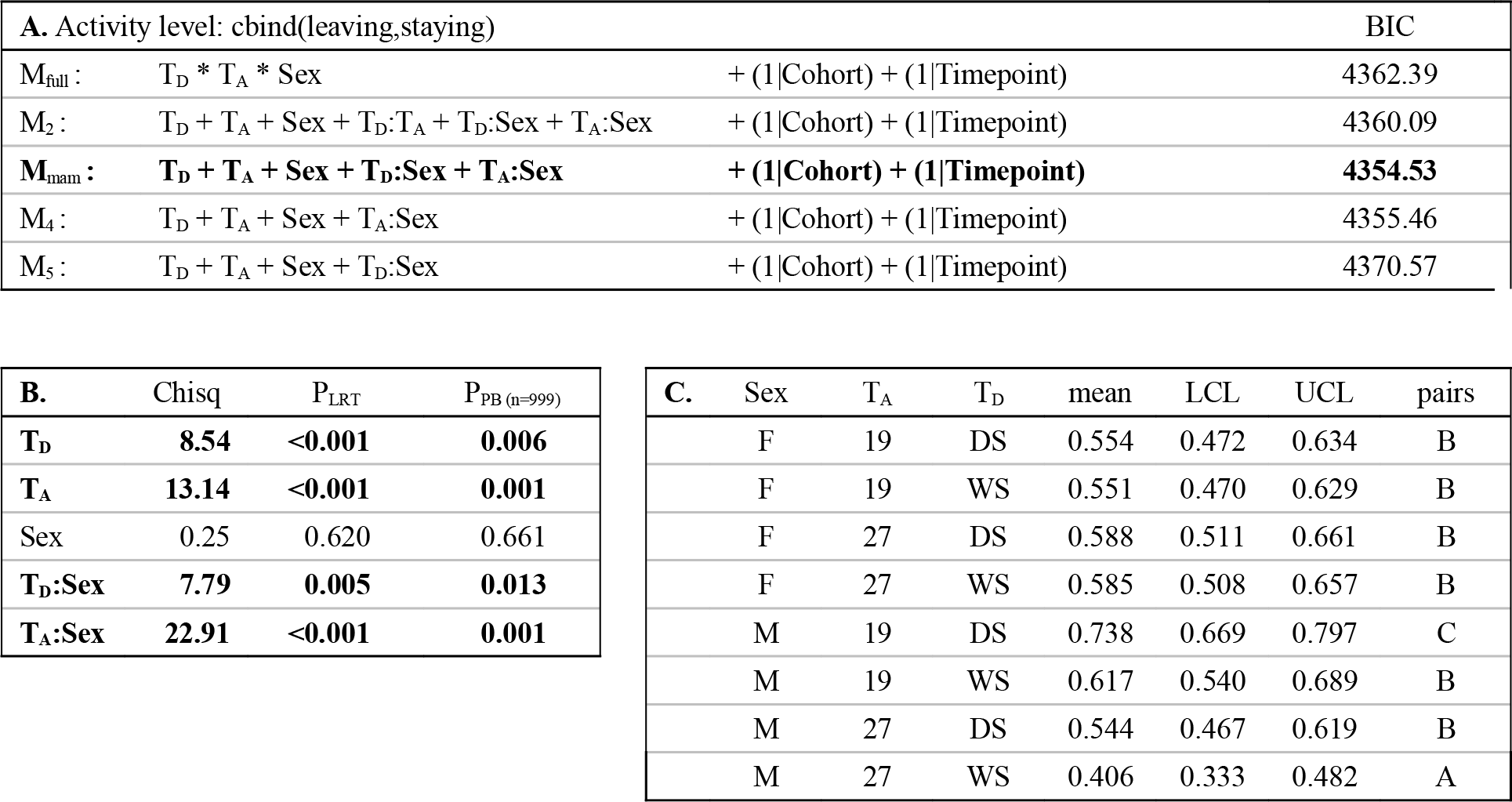
GLMM for activity levels (related to Fig 2b in the main text). **A.** Minimum adequate models were obtained by backward elimination, starting from the full model and using Bayesian Information Criterion (BIC). **B.** P-values for the minimum adequate model were calculated via likelihood ratio tests and parametric bootstrap (nsim=999), using the package *afex*. **C.** Mean probabilities, as well as upper and lower confidence limits, are given for each sex and experimental treatment. Post-hoc pairwise comparisons (alpha=0.05), obtained using the package *lsmeans*, specify differences between experimental treatments and sexes.

